# Knowledge, Attitude and Practice of Breast Cancer Screening among women attending primary care centers in Abu Dhabi

**DOI:** 10.1101/652552

**Authors:** Noora Ali Al Blooshi, Ruqayya Saaed Al Mazrouei, Hind Nasser Al Razooqi, Ebtihal Ahmad Darwish, Maha Mohamed Al Fahim, Fozia Bano Khan

## Abstract

**Background:** Breast cancer is the most common cancer among women in UAE. Screening for it can reduce morbidity and mortality and improve women survival. Low level of knowledge and poor practice of breast cancer screening could be due to many factors. The aim of our study is to assess the knowledge, attitude and practice of breast cancer screening of women attending primary care centers in Abu Dhabi region.

**Method:** A cross sectional study was done in 2017 using questionnaire about breast cancer awareness measure. Six primary health care centers were included which were located inside and outside Abu Dhabi island. Three hundred eighty three women participated in the study, between the age of 40-65.

**Results:** The facilities to screen for breast cancer screening is available, but it is still underutilized by women. Even though we found in our study that women had high level of knowledge about breast cancer (45.7%), but their practice for mammogram was poor (52.2%). We also found that, the higher the education, employment and family history of breast cancer women had better knowledge, with statistically significant result respectively (P=0.000), (P=0.018), (P=0.013), and women older than 49 had better practice of mammogram (P=0.000). In our study, we concluded that women who attend clinic located inside island of Abu Dhabi had better knowledge and practice compared to women attending clinics located outside the island who had better attitude.

**Conclusion:** In our study we found that despite having the modalities and services for breast cancer screening, it is still underutilized. Our population sample showed that women had good knowledge about breast cancer but poor practice for mammogram. Educational level, age and region all played role in their knowledge, attitude and practice. As primary care providers we are a big influencers on the society and the first contact to them, we can utilize this to spread the awareness. We can approach the women through social media, more campaigns and offering free mammogram to women who cannot afford paying for it. Spreading the awareness about screening will reduce the burden of breast cancer treatment on the health care system government too.

## Background

Breast cancer is a major health burden worldwide, impacting 2.1 million women annually and also a leading cause of death among women worldwide, with an estimate of 1.7 million cases and 521,900 deaths in 2012 [1]. According to the International Agency for Research on Cancer, it now represents one in four of all cancers in women.

Globally, breast cancer incidence and mortality are expected to increase by 50% between 2002 and 2020. The highest incidence remains in the developed world, however incidence rates are increasing in other areas of the world [4]. The incidence rates vary globally ranging from 91.6 per 100,000 women in North America to 43 per 100,000 in the Middle East and Northern Africa (MENA) region (International Agency for Research on Cancer, 2012).

Breast cancer remains a major public health threat among women in the Arab world, with the United Arab Emirates (UAE) ranking the seventh country with a continuous increase in the incidence rate. According to the UAE Cancer Registry 2014, it was the leading cause of death with a total of 81 deaths in the year 2014 [7]. It mostly affects Arab women at younger ages compared to women in western countries (WHO). A recent study in 2014 showed that breast cancer was the second leading cause of death among women in Abu Dhabi, comprising 20.3% of the five most common cancers in 2014 [2,3,6].

The impact of breast cancer can be reduced through various practices including breast self-examination (BSE), clinical breast examination (CBE), or obtaining a mammogram. The UAE government has made great efforts in promoting early cancer detection through various health authorities. The Health Authority of Abu Dhabi established a breast cancer screening program since 2008. Despite this, screening rates have not yet reached levels of breast practice.

A recent study done in 2014 in Al Ain UAE, showed that lack of knowledge about breast cancer is not the only reason for the delay in early presentation. Other factors such as personal, social and cultural factors also contribute. Furthermore, a study in 2010, conducted in Dubai UAE, showed a low level of practice of breast cancer screening among local women, in addition to the unawareness of the common risk factors and the signs and symptoms of breast cancer [3].

No other studies have been conducted to assess the impact of breast cancer screening in Abu Dhabi. Therefore our study aims to measure the level of knowledge, attitude, and practice of breast cancer screening in women attending primary health care centers in Abu Dhabi.

## Method

### Study design and participant

This cross sectional study was conducted among women visiting six primary healthcare clinics, AlBateen, Zafaranah, Rowda, Maqta, Mohammed Bin Zayed and Baniyas, in Abu Dhabi between August 2016 till August 2017 using a self-administered questionnaire.

The target population for the study were women between the age of 40-65 years old who attended primary care centers within 40km from the Abu Dhabi Island and regardless their nationality. The selection was stratified random selection and involved who agreed to participate in the study. We excluded women who were non Abu Dhabi resident, non-English or non-Arabic speakers and women who had personal history of breast cancer. The estimated target population size was 168,286 women between the ages of 40 and 65 were identified based on the statistical yearbook of Abu Dhabi 2015. The sample size was calculated using a 95% confidence interval and 5% margin of error to be 383 participants.

The self-administered questionnaires were then proportionally distributed to the six primary health care clinics based on the number of patients attending each clinic from the total sample size.

At each clinic, questionnaires were distributed to women who met inclusion and exclusion criteria in a stratified random collection sampling method until the required numbers were attained.

### Study instrument and ethical approval

An Arabic and English versions questionnaires were developed and used in the present study which was adapted from a similar studies that were conducted in Al Ain City [8], RAK city [10] and Saudi Arabia [9] and modified to meet our objectives. The self-administered questionnaire contained a total of 102 questions and was divided into four parts that addressed the participants’ sociodemographic characteristics (17 questions) and breast cancer screening knowledge (28 questions), attitudes (5 question), and practices (7 questions). A scale scoring system was used to categorize knowledge, attitude and practice as follow: Knowledge: Good (19–27 points), Fair (9–18 points), Poor (0–8 points); Attitude: positive (4 points), neutral (2-3 points), negative (0–1 point); Practice: Good (2 points), bad (0–1 point).

Informed consent forms were attached with each questionnaire for the participants to read and sign if they were willing to participate in the study. Questionnaires and consent forms were drafted in English and Arabic versions. A pilot study was conducted after ethical approval was granted in order to assess the questionnaire’s comprehensibility, and modifications were accordingly made.

### Data Collection

Questionnaires were printed and then proportionally distributed to the six primary healthcare clinics in Abu Dhabi. The charge nurses at these health center were briefed regarding the questionnaire and given training regarding the participants’ anonymity and informed consent.

The charge nurse at each clinic was requested to distribute the questionnaires randomly to patients who matched our inclusion criteria. After the completion of the questionnaires by the participants, they collected and sealed the questionnaires in envelopes to ensure the participants’ confidentiality. After the end of the study period, the charge nurse was requested to return the completed questionnaires to the authors.

### Data Analysis

After the collection of the questionnaires, the obtained data were organized using the MS Excel software program, coded, and analyzed using the Statistical Package for Social Sciences (SPSS) version 18. Means and standard deviation (SD) were used for numerical data, whereas percentages were used for categorical data.

First, chi squared (χ2) test was conducted to assess the effect of certain factors on breast cancer screening knowledge, attitudes, and practices. Factors that were analyzed in this study included: Age, education, employment status, marital status, monthly income, menopause history, age of menopause, family history of breast cancer, age of first pregnancy, living children, breast feed their child.

## Results

A total of 383 questionnaires were distributed.

### Description and characteristics of participants

The majority of the participant were women aged 40-48 years old (58.7%), local (54.2%), and married (75.5%). Among the participants 45% had university and above education, 55% were unemployed and 70% had sufficient monthly income (Table 1).

**Table 1.** Socio-Demographic characteristics of study sample

| **Table 1. Socio- Demographic characteristics of study sample** |  |  |
| --- | --- | --- |
|  | Number | Percent (%) |
| **Age (n=378 )** |  |  |
| 40-48 | 222 | 58.7 |
| 49-58 | 113 | 29.9 |
| 59-65 | 43 | 11.4 |
| **Nationality (n=378 )** |  |  |
| Local | 205 | 54.2 |
| Non local | 173 | 45.8 |
| **Education (n=380 )** |  |  |
| Primary and less | 94 | 24.7 |
| Secondary | 115 | 30.3 |
| University and above | 171 | 45.0 |
| **Employment status (n=371 )** |  |  |
| Employed | 167 | 45.0 |
| Unemployed | 204 | 55.0 |
| **Marital status (n=371 )** |  |  |
| Single | 35 | 9.20 |
| Married | 287 | 75.5 |
| Divorce/widow | 58 | 15.3 |
| **Monthly income (n=357 )** |  |  |
| Sufficient | 250 | 70.0 |
| Sufficient and saving | 45 | 12.6 |
| Insufficient | 62 | 17.4 |

### Women’s Breast cancer screening knowledge

The table contain the correctly answered questioned regarding knowledge of breast cancer screening among women attending primary care centers in Abu Dhabi.

A total of (45.7%) found to have good knowledge about breast cancer screening, (48.8%) had fair knowledge, and only (5.5%) had poor knowledge. 82.2% correctly reported that breast cancer is the most common cancer. Regarding risk factors for breast cancer, 91.6% reported that breast feeding protect against breast cancer, 84.6% reported that it is not related to the size of the breast, 79.4% reported that having first degree relative with breast cancer increase risk of own breast cancer, and 62.9% reported that it is not related to age.

Regarding symptoms of breast cancer, 79.6 % reported painless mass could present, 50.9% reported that bloody nipple discharge, and 43.1% reported nipple retraction as symptoms.

For the breast cancer screening method, 75.5% reported mammogram as being a screening method, 67.6% reported breast self-examination, and 50.9% reported clinical breast examination.

78.1%, 57.2% and 32.4% of women reported that all women above 40, women who have family history of breast cancer, and who had personal history of breast cancer must do regular mammogram respectively.

The most common sources of information regarding breast cancer screening were from primary health care providers (41.0%), TV or Radio (31.1%), Magazine of Brochure (31.1%) and Hospital Doctors (27.2%) (Table 2).

**Table 2.** Breast Cancer screening Knowledge of women visiting primary care centers in Abu Dhabi (n=383)

|  |  |  |
| --- | --- | --- |
|  | Number of correct answer | (%) of correct answer |
| Breast cancer most common cancer among women | 371 | (82.2) |
| **Risk Factors of breast cancer** |  |  |
| Breast cancer not related to age | 241 | 62.9 |
| Breast cancer not related to size of breast | 324 | 84.6 |
| Having first degree relative increase risk of breast cancer | 304 | 79.4 |
| Stress is not a risk factor for breast cancer | 208 | 54.3 |
| Using HRT increase risk of breast cancer | 154 | 40.2 |
| Breast feeding protect against breast cancer | 351 | 91.6 |
| Early Menarche increase risk of breast cancer | 38 | 9.90 |
| **Symptoms of breast cancer** |  |  |
| Bloody discharge for the nipple | 195 | 50.9 |
| Painless mass | 305 | 79.6 |
| Pelvic pain is not a symptom | 328 | 85.6 |
| Absence of period is not a symptom | 313 | 81.7 |
| Nipple retraction | 165 | 43.1 |
| **Breast cancer screening methods** |  |  |
| Breast self-examination | 259 | 67.6 |
| Clinical Breast examination | 195 | 50.9 |
| Mammogram | 289 | 75.5 |
| US is not screening method | 268 | 70.0 |
| MRI is not screening method | 330 | 86.2 |
| CT is not screening method | 327 | 85.4 |
| Frequency of Breast self-examination is once a month | 161 | 42.0 |
| Right time to do BSE is end of menses | 190 | 49.6 |
| Mammogram can detect breast cancer early | 312 | 81.5 |
| Frequency of Mammogram is every 2 years | 155 | 40.5 |
| **Group that must do regular mammogram** |  |  |
| All women above 40 | 299 | 78.1 |
| Women who have family history of breast cancer | 219 | 57.2 |
| Personal history of breast cancer | 124 | 32.4 |
| Female above 18 should not do regular mammogram | 314 | 82.0 |
| **Participant overall knowledge** |  |  |
| Good | 175 | 45.7 |
| Fair | 187 | 48.8 |
| Poor | 21 | 5.5 |
| **Participant source of information about breast cancer screening** |  |  |
| TV or Radio | 119 | 31.1 |
| Magazine or Brochure | 119 | 31.1 |
| Hospital Doctor | 104 | 27.2 |
| Primary Health care Provider | 157 | 41 |
| Others | 56 | 14.6 |

### Factors Affecting Women’s Breast Cancer Screening knowledge

Table 3 shows that as the level of education increase, the level of knowledge regarding breast cancer improve. This results is statistically significant (P=0.00). Better breast cancer screening knowledge was found among women who were employed (P = 0.018), and those who had family history of breast cancer (P = 0.013). Women who had Good and Fair knowledge found to have a better attitude and practice regarding breast cancer screening than those with poor knowledge (P = 0.000) (P = 0.025) respectively.

**Table 3.**
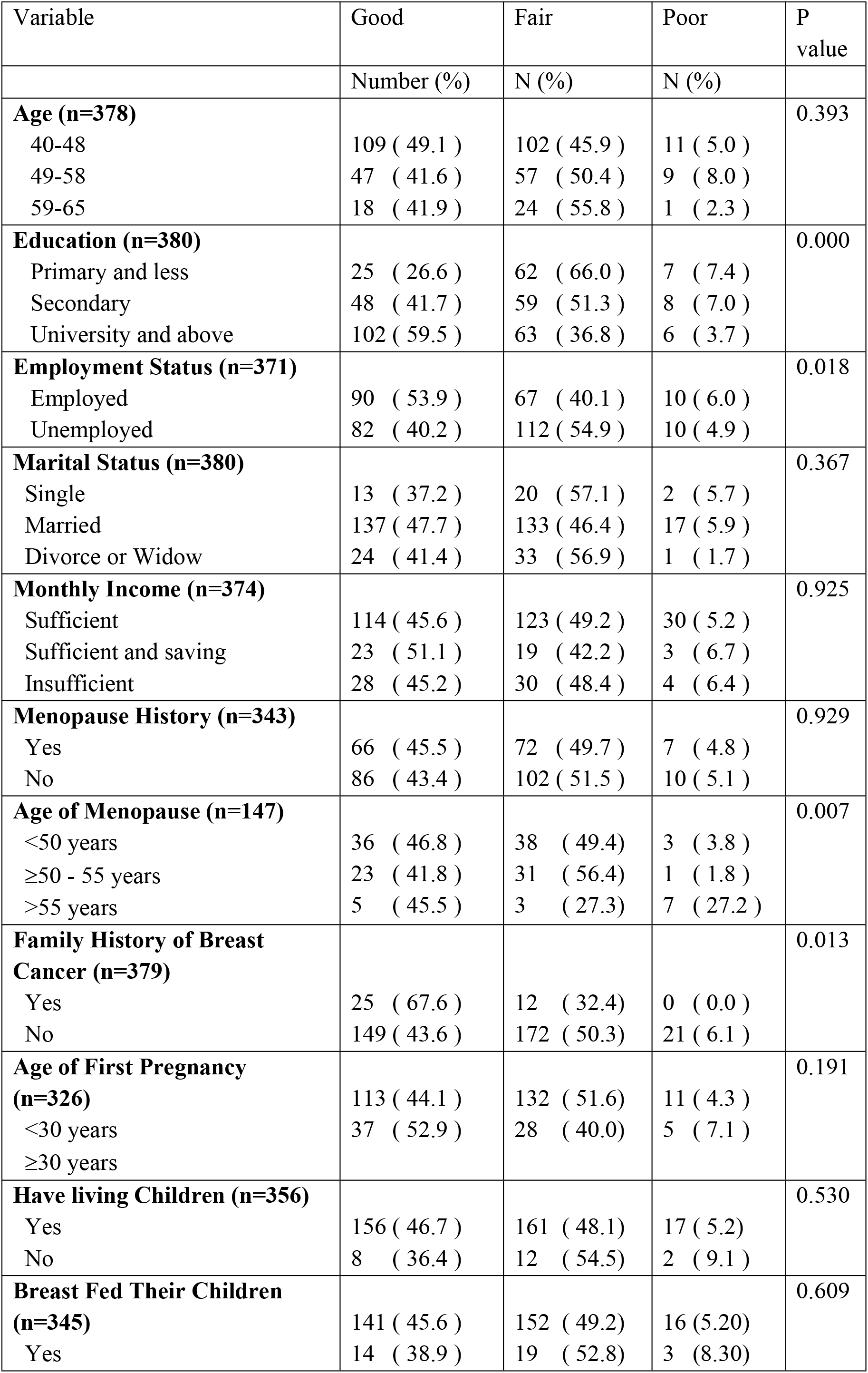
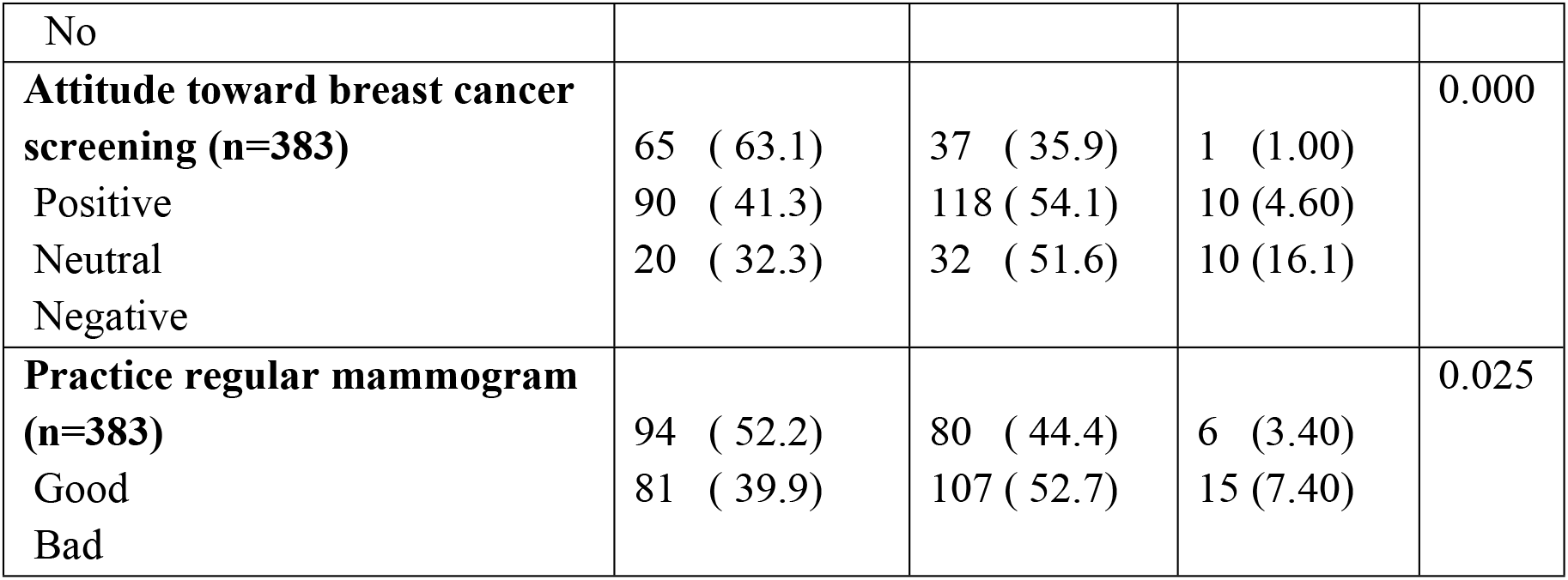
Factors Affecting Women’s Breast Cancer Screening Knowledge

There was no statistically significant relationship between level of knowledge and age (P = 0.393), Marital Status (P = 0.367), Monthly income (P = 0.925), Age of First Pregnancy (P = 0.191), Have living Children (P = 0.530) and Breast Fed Their Children (P = 0.609).

### Women Breast Cancer screening attitude

The table contain the correctly answered questioned regarding attitude of breast cancer screening among women attending primary care centers in Abu Dhabi.

A total of (26.9%) found to have positive attitude about breast cancer screening, (56.9%) had neutral attitude, and only (16.2%) had Negative attitude (Table 4).

**Table 4.** Breast Cancer screening Attitude of women visiting primary care centers in Abu Dhabi (n=383)

| **Table 4** |  |  |
| --- | --- | --- |
|  | Number of correct answer | % of correct answer |
| **Important for early detection** | 349 | 91.1 |
| **Painful procedure** | 101 | 30.6 |
| **Done regularly** | 302 | 78.9 |
| **Easy to perform** | 234 | 74.1 |
| **Increase anxiety** | 144 | 37.6 |
| **Overall altitude** |  |  |
| Positive | 103 | 26.9 |
| Neutral | 218 | 56.9 |
| Negative | 62 | 16.2 |

### Factors Affecting Women’s Breast Cancer Screening Attitude

Table 5 shows that women among older population had a better attitude toward breast cancer screening. This results is statistically significant (P=0.016).

**Table 5.** Factors Affecting Women’s Breast Cancer Screening Attitude

| **Table 5. Factors Affecting Women's Breast Cancer Screening Attitude** |  |  |  |  |
| --- | --- | --- | --- | --- |
| Variable | Positive | Neutral | Negative | P value |
|  | Number (%) | Number (%) | Number (%) |  |
| **Age (n=378)** |  |  |  | 0.016 |
| 40-48 | 61 ( 27.5 ) | 118 ( 53.2 ) | 43 ( 19.3 ) |  |
| 49-58 | 24 ( 21.2 ) | 75 ( 66.4 ) | 14 ( 12.4 ) |  |
| 59-65 | 18 ( 41.9 ) | 22 ( 51.1 ) | 3 ( 7.0 ) |  |
| **Education (n=380)** |  |  |  | 0.591 |
| Primary and less | 28 ( 29.8 ) | 56 ( 59.6 ) | 10 ( 10.6 ) |  |
| Secondary | 30 ( 26.1 ) | 64 ( 55.7 ) | 21 ( 18.2 ) |  |
| University and above | 45 ( 26.3 ) | 96 ( 56.1 ) | 30 ( 17.6 ) |  |
| **Employment Status (n=371)** |  |  |  | 0.017 |
| Employed | 39 ( 23.4 ) | 93 ( 55.6 ) | 35 ( 21.0 ) |  |
| Unemployed | 63 ( 30.9 ) | 119 ( 58.3 ) | 22 ( 10.8 ) |  |
| **Marital Status (n=344)** |  |  |  | 0.540 |
| Single | 7 ( 20.0 ) | 20 ( 57.1 ) | 8 ( 22.9 ) |  |
| Married | 79 ( 27.5 ) | 126 ( 56.4 ) | 46 ( 16.1 ) |  |
| Divorce or Widow | 17 ( 29.3 ) | 35 ( 60.3 ) | 6 ( 10.4 ) |  |
| **Monthly Income (n=357)** |  |  |  | 0.990 |
| Sufficient | 70 ( 28.0 ) | 141 ( 56.4 ) | 39 ( 15.6 ) |  |
| Sufficient and saving | 13 ( 28.9 ) | 26 ( 57.8 ) | 6 ( 13.3 ) |  |
| Insufficient | 16 ( 25.8 ) | 36 ( 58.1 ) | 10 ( 16.1 ) |  |
| **Menopause History (n=343)** |  |  |  | 0.437 |
|  | 35 ( 24.1 ) | 91 ( 62.8 ) | 19 ( 13.1 ) |  |
| Yes | 54 ( 27.3 ) | 111 ( 56.1 ) | 33 ( 16.6 ) |  |
| No |  |  |  |  |
| **Age of Menopause (n=145)** |  |  |  | 0.299 |
| <50 years | 19 ( 24.7 ) | 48 ( 62.3 ) | 10 ( 13.0 ) |  |
| ≥50 - 55 years | 17 ( 30.9 ) | 30 ( 54.5 ) | 8 ( 14.6 ) |  |
| >55 years | 0 ( 00.0 ) | 9 ( 81.8 ) | 2 ( 18.2 ) |  |
| **Family History of Breast Cancer (n=379)** |  |  |  | 0.180 |
| Yes | 12 ( 32.4 ) | 23 ( 62.2 ) | 2 ( 5.40 ) |  |
| No | 91 ( 26.6 ) | 193 ( 56.4 ) | 58 ( 17.0 ) |  |
| **Age of First Pregnancy (n=326)** |  |  |  | 0.528 |
| <30 years | 80 ( 31.2 ) | 139 ( 54.3 ) | 37 ( 14.5 ) |  |
| ≥30 years | 17 ( 24.3 ) | 42 ( 60.0 ) | 11 ( 15.7 ) |  |
| **Have living Children (n=356)** |  |  |  | 0.041 |
| Yes | 98 ( 29.3 ) | 185 ( 55.4 ) | 51 ( 15.3 ) |  |
| No | 1 ( 4.5 ) | 16 ( 72.7 ) | 5 ( 22.8 ) |  |
| **Breast Fed Their Children (n=345)** |  |  |  | 0.039 |
| Yes | 93 ( 30.1 ) | 169 ( 54.7 ) | 47 ( 15.2 ) |  |
| No | 4 ( 11.1 ) | 27 ( 75.0 ) | 5 ( 13.9 ) |  |
| **Practice (n=383)** |  |  |  | 0.000 |
| Good (practiced mammogram) | 63 (35.0) | 99 (55.0) | 18(10.0) |  |
| Bad (did not practice mammogram) | 40 (19.7) | 119 (58.6) | 44(21.7) |  |

Positive breast cancer screening attitude was found among women who were unemployed (P = 0.017), who have living children (P = 0.041), and women who breast fed their children (P = 0.039).

Women who had positive and Neutral attitude found to have a better practice regarding breast cancer screening than those with negative attitude (P = 0.000). There was no statistically significant relationship between attitude and education (P = 0.591), Marital Status (P = 0.540), Monthly income (P = 0.990), Menopause history (P = 0.437), Age of menopause (P = 0.299), Family history of breast cancer (P = 0.180), Age of First Pregnancy (P = 0.528).

### Women Breast Cancer screening Practice

The table contain the correctly answered questioned regarding practice of breast cancer screening among women attending primary care centers in Abu Dhabi.

A total of (47%) found to have Good practice about breast cancer screening by using mammogram, (53%) had Bad practice.

59.7% practiced Breast self-examination, 41.0% was doing it regularly and 24.7% was doing it monthly. 35.0%, 26.4%, and 20.0% of the women reported that the reasons for not practicing were lack of knowledge, don’t know how to do it, and had fear to find something abnormal.

48.9% among the women practiced clinical breast examination and 47.0% was doing it once a year.

64.5% among the women practiced mammogram and 78.3% was practicing it once 1-3 years (Table 6).

**Table 6.**
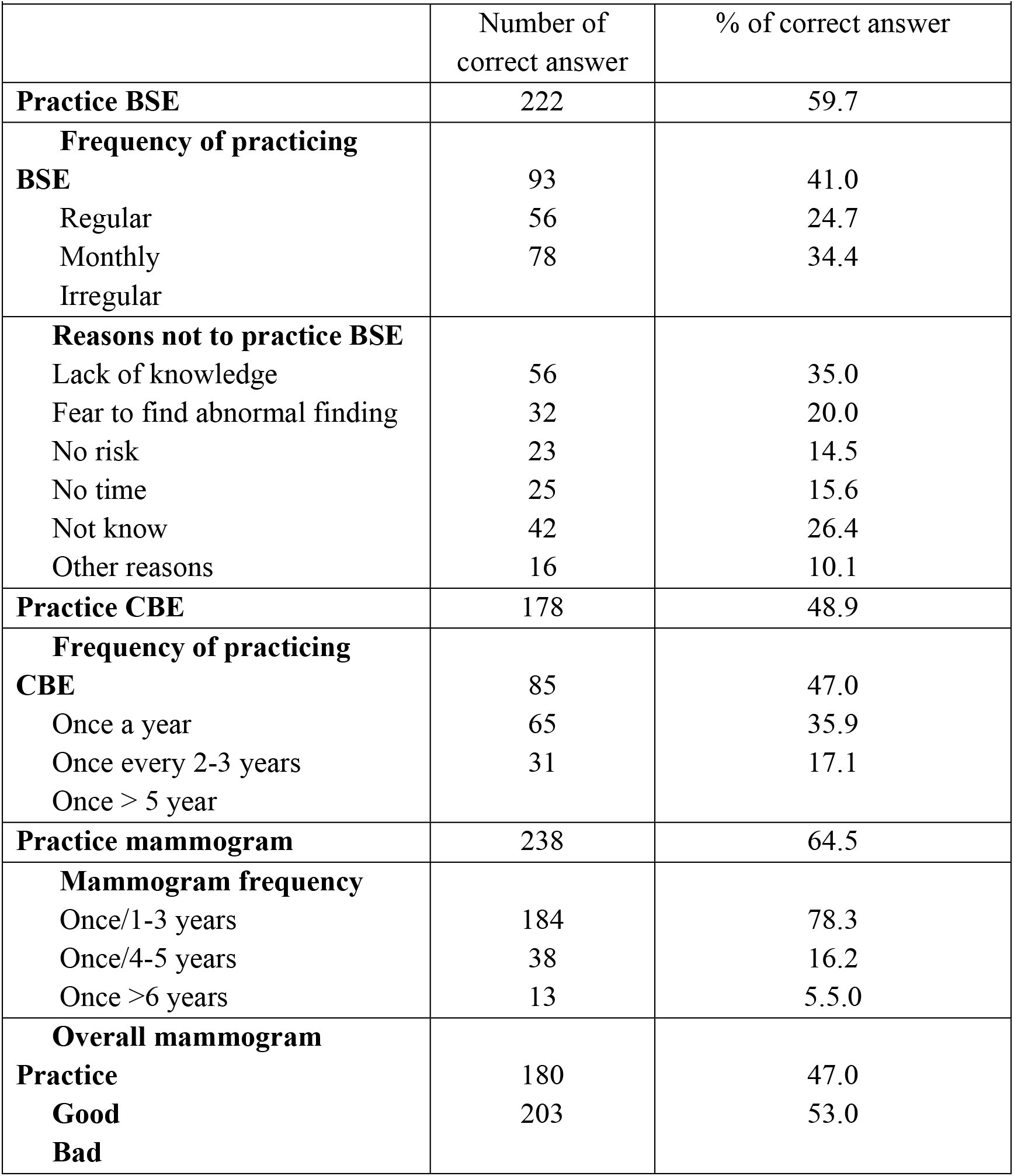
Breast Cancer screening Practice of women visiting primary care centers in Abu Dhabi (n=383)

### Factors Affecting Women’s Breast Cancer Screening Practice

Table 7 shows that women among older population had a good practice of mammogram. This results is statistically significant (P=0.00).

**Table 7.** Factors Affecting Women’s Breast Cancer Screening Practice

| **Table 7. Factors Affecting Women's Breast Cancer Screening Practice** |  |  |  |
| --- | --- | --- | --- |
| Variable | Good (practiced mammogram) | Bad (did not practice mammogram) | P value |
|  | Number (%) | Number (%) |  |
| **Age (n=378)** |  |  | 0.000 |
| 40-48 | 83 ( 37.4 ) | 139( 62.6 ) |  |
| 49-58 | 71 ( 62.8 ) | 42 ( 37.2 ) |  |
| 59-65 | 25 ( 58.1 ) | 18 ( 41.9 ) |  |
| **Education (n=380)** |  |  | 0.728 |
| Primary and less | 47 ( 50.0 ) | 47 ( 50.0 ) |  |
| Secondary | 55 ( 47.8 ) | 60 ( 52.2 ) |  |
| University and above | 77 ( 45.0 ) | 94 ( 55.0 ) |  |
| **Employment Status (n=371)** |  |  | 0.377 |
| Employed | 75 ( 44.9 ) | 92 ( 55.1 ) |  |
| Unemployed | 101 ( 49.5 ) | 103 ( 50.5 ) |  |
| **Marital Status (n=380)** |  |  | 0.006 |
| Single | 10 ( 28.6 ) | 25 ( 71.4 ) |  |
| Married | 133 ( 46.3 ) | 154 ( 53.7 ) |  |
| Divorce or Widow | 36 ( 62.1 ) | 22 ( 37.9 ) |  |
| **Monthly Income (n=357)** |  |  | 0.674 |
| Sufficient | 114 ( 45.6 ) | 136 ( 54.4 ) |  |
| Sufficient and saving | 22 ( 48.9 ) | 23 ( 51.1 ) |  |
| Insufficient | 32 ( 51.6 ) | 30 ( 48.4 ) |  |
| **Menopause History (n=343)** |  |  | 0.000 |
| Yes | 84 ( 57.9 ) | 61 ( 42.1 ) |  |
| No | 77 ( 38.9 ) | 121 ( 61.1 ) |  |
| **Age of Menopause (n=143)** |  |  | 0.292 |
| <50 years | 43 ( 55.8 ) | 34 ( 44.2 ) |  |
| ≥50 - 55 years | 34 ( 61.8 ) | 21 ( 38.2 ) |  |
| >55 years | 4 ( 36.4 ) | 7 ( 63.6 ) |  |
| **Family History of Breast Cancer (n=379)** |  |  | 0.024 |
| Yes | 24 ( 64.9 ) | 13 ( 35.1 ) |  |
| No | 155( 45.3 ) | 187 ( 54.7) |  |
| **Age of First Pregnancy**<br>**(n=326)** |  |  | 0.744 |
| <30 years | 119 ( 46.5 ) | 137 (53.5 ) |  |
| ≥30 years | 31 ( 44.3 ) | 39 ( 55.7 ) |  |
| **Have living Children**<br>**(n=356)** |  |  | 0.129 |
| Yes | 162 ( 48.5 ) | 172 ( 51.5) |  |
| No | 7 ( 31.8 ) | 15 ( 68.2 ) |  |
| **Breast Fed Their Children**<br>**(n=345)** |  |  | 0.783 |
| Yes | 147 ( 47.6 ) | 162 ( 52.4 ) |  |
| No | 18 ( 50.0 ) | 18 ( 50.0 ) |  |

Good breast cancer screening practice was found among women who were married (P = 0.006), with menopause history (P = 0.000), and those who had family history of breast cancer (P = 0.024).

There was no statistically significant relationship between practice and education (P = 0.728), Employment status (P = 0.377), Monthly income (P = 0.674), Age of menopause (P = 0.292), Age of First Pregnancy (P = 0.744), Having living children (P = 0.129) and breast Fed their children (P = 0.783).

### Overall knowledge, attitude and practice between primary care clinics in Abu Dhabi

While comparing between clinics within and outside Abu Dhabi island, best level of knowledge among women attending the clinics and best breast cancer screening practice were found in Rowdah followed by Zafrana and followed by AlBateen. However, the most positive attitude among women attending primary care centers were found in Baniyas, followed by Al Bateen and followed by Maqta. These results were statistically significant (P=0.000, P=0.003 and P=0.000) respectively (Table 8).

**Table 8.** Overall knowledge, attitude and practice between primary care clinics in Abu Dhabi

|  |  |  |  |  |  |  |  |
| --- | --- | --- | --- | --- | --- | --- | --- |
|  | **Al Bateen** | **Rowdah** | **Zafrana** | **Maqta** | **MBZ** | **Baniyas** | **P** |
|  | NUMBER<br>(%) | NUMBER<br>(%) | NUMBER<br>(%) | NUMBER<br>(%) | NUMBER<br>(%) | NUMBER<br>(%) | Value |
| **Overall Knowledge** |  |  |  |  |  |  | 0.000 |
| Good | 27 (47.3) | 29 (63.0) | 48 (50.5) | 20 (37.0) | 15 (28.3) | 36 (46.2) |  |
| Fair | 29 (50.9) | 16 (34.8) | 38 (40.0) | 34 (63.0) | 30 (56.6) | 40 (51.2) |  |
| Poor | 1 (1.80) | 1 (2.20) | 9 (9.50) | 0 (0.00) | 8 (15.1) | 2 (2.60) |  |
| **Overall Attitude** |  |  |  |  |  |  | 0.000 |
| Positive | 20 (35.1) | 10 (21.7) | 16 (16.8) | 14 (25.9) | 5 (9.40) | 38 (48.7) |  |
| Neutral | 28 (49.1) | 30 (65.2) | 58 (61.1) | 30 (55.6) | 34 (64.2) | 38 (48.7) |  |
| Negative | 9 (15.8) | 6 (13.1) | 21 (22.1) | 10 (18.5) | 14 (26.4) | 2 (2.60) |  |
| **Overall mammogram**<br>**Practice** |  |  |  |  |  |  | 0.003 |
| Good | 29 (50.9) | 30 (65.2) | 52 (53.7) | 26 (48.1) | 17 (32.1) | 27 (34.6) |  |
| Bad | 28 (49.1) | 16 (34.8) | 44 (46.3) | 28 (51.9) | 36 (67.9) | 51 (65.4) |  |

## Discussion

Breast cancer is the most common cancer among females in UAE according to a 2011 report released by the department of health in Abu Dhabi. As primary health care providers we are the first contact for providing health care services, such as prevention services and early detection of breast cancer. Our study compared differences between Knowledge, attitude and practice of breast cancer screening among women in Abu Dhabi region, and the results of other studies conducted in Ras A Khaimah, Al Ain and Najran, Saudi Arabia studies.

While comparing the level of knowledge about breast cancer in our study with Saudi Arabia, and Al Ain studies, we found that 45.7%, 10.2%, and 5% of the women had good knowledge respectively. The highest level of our percentage could be attributed that most women included could be at younger age and have access to evidence based medicine while searching the internet. Another reason could be women are being influenced by people in social media or they are becoming more trust worthy with their primary care providers regarding health related problems.

Surprisingly we found similar outcome in Al Ain study regarding knowledge about risk factors of breast cancer. Our results showed that women knows that breast cancer is the most common cancer among women in UAE, breast feeding is protective against breast cancer but they were not sure when mammogram should be done.

Both studies showed that the younger the population and the higher the education level the better the knowledge.

Like we concluded above, social media plays an important rule likewise the internet. And more campaigns are done in universities, schools and in the communities spreading the awareness about the importance of breast cancer screening.

Our research and Saudi Arabia research showed that with higher education level, being employed and having a personal or family history of breast cancer had a good impact on the knowledge about breast cancer screening. The reason behind that could be the primary care doctors are providing good consultation in their visits. This is reflected in our research which showed that primary health care providers were the primary source for the women in Abu Dhabi. Social media also plays a rule nowadays for spreading news about importance of breast cancer screening. In contrast, Saudi Arabia research showed that most women get their knowledge from social media and only 8.8% received there information from primary care doctor. Furthermore, in Al Ain study showed that 38% of women got their information form heath care provider.

When comparing between inside and outside the island of Abu Dhabi, best knowledge was found among women living inside the island. The reason behind that could be that more campaigns about cancer are happened inside compared to outside the island. Women who live outside the island (rural place) are more likely to be house wives and somehow they have low level of education.

The study showed regardless of having free service of mammogram among women, there is lack of knowledge and underutilization of the service. Same results were found in the studies done in Al Ain and in Saudi Arabia.

In our study it was shown that 47% of women practiced mammogram, compared to 44.9% in Al Ain study, 37.6% in Ras Al Khaimah, and 15% in Saudi Arabia study. Even though we had the highest rate of mammogram practice, overall it is still considered poor practice. Although our study showed that women in Abu Dhabi have good knowledge about breast cancer screening, they still have poor practice. Reason could be that some women does not feel comfortable to be examined by doctors, they only seek medical advice when they discover abnormal finding in their breasts, too busy in their social, occupational and personal life, or don’t have enough time to get general checkup.

In our study women who practiced mammogram were more compared to other studies. This could be attributed to the awareness campaigns which are conducted by the UAE health authorities, and offering free mammograms in Oct-Nov period (Breast cancer awareness month) for women with insurance not covering mammogram.

Both our study and the one conducted in Al Ain showed that older women ≥ 49 years old more likely to go for mammogram. Reasons behind that could be younger population women <49 think they are not the target for the screening program. Older women are having chronic disease that they seek medical advice and their primary health care providers are offering mammogram more often than to younger population who only seek their doctors when they are ill.

When comparing between clinics, allocation plays an important rule. The ones which were located inside the island had good practice of mammogram compared to outside the island. The reasons might be that, women inside the island tend to be more knowledgeable and aware about the importance of breast cancer screening. Moreover, more campaigns are done inside Abu Dhabi island compared to outside.

## Conclusion

In our study we found that despite having the modalities and services for breast cancer screening, it is still underutilized. Our population sample showed that women had good knowledge about breast cancer but poor practice for mammogram. Educational level, age and region all played role in their knowledge, attitude and practice. As primary care providers we are a big influencers on the society and the first contact to them, we can utilize this to spread the awareness. We can approach the women through social media, more campaigns and offering free mammogram to women who cannot afford paying for it. Spreading the awareness about screening will reduce the burden of breast cancer treatment on the health care system government too.

## Acknowledgement

We would like to express our deep gratitude to Milany, Baverleen, Heba, Shadia and Khaleel the charge nurses of the primary care centers, for their hard work and patience. For helping with distributing questionnaire and keeping it sealed and private.

